# T cell responsiveness to IL-10 defines the immunomodulatory effect of costimulation blockade via anti-CD154 and impacts transplant survival

**DOI:** 10.1101/2024.06.12.598652

**Authors:** Marcos Iglesias, Darrel Bibicheff, Alexander Komin, Maria Chicco, Samantha Guinn, Brendan Foley, Giorgio Raimondi

**Author notes:** **Corresponding authors e-mail**: Giorgio Raimondi and Marcos Iglesias.

## Abstract

Costimulation blockade (CoB)-based immunotherapy is a promising alternative to immunosuppression for transplant recipients; however, the current limited understanding of the factors that impact its efficacy restrains its clinical applicability. In this context, pro- and anti-inflammatory cytokines are being recognized as having an impact on T cell activation beyond effector differentiation. This study aims at elucidating the impact of direct IL-10 signaling in T cells on CoB outcomes. We used a full-mismatch skin transplantation model where recipients had a T cell-restricted expression of a dominant negative IL-10 receptor (10R-DN), alongside anti-CD154 as CoB therapy. Unlike wild-type recipients, 10R-DN mice failed to benefit from CoB. This accelerated graft rejection correlated with increased accumulation of T cells producing TNF-α, IFN-γ, and IL-17. In vitro experiments indicated that while lack of IL-10 signaling did not change the ability of anti-CD154 to modulate alloreactive T cell proliferation, the absence of this pathway heightened T_H_1 effector cell differentiation. Furthermore, deficiency of IL-10 signaling in T cells impaired Treg induction, a hallmark of anti-CD154 therapy. Overall, these findings unveil an important and novel role of IL-10 signaling in T cells that defines the success of CoB therapies and identifies a target pathway for obtaining robust immunoregulation.

## 1. Introduction

Induction of transplant tolerance via regimens that do not require dangerous conditioning or that do not limit the pool of eligible patients remains elusive. A major effort is currently on the optimization of “costimulation blockade” (CoB) therapies, where blockade of activatory signals of alloreactive T cells have proved to be a potent inhibitor of rejection in multiple rodent models.^1,2^ The cellular mechanisms identified as responsible for this effect are a combination of induction of hyporesponsive alloreactive T-cells (anergy), deletion, and formation of donor-specific regulatory T cells (Tregs).^3,4^ Implementation of this strategy in the clinic however has been difficult.^5^ The identification and control of the factors and mechanisms that affect the regimen’s efficacy is essential to overcome the observed limitations and improve clinical outcomes.

The presence of infections or inflammatory cues at the time of transplantation are among the main barriers that prevent the induction of tolerance with CoB therapies.^6–8^ The mechanisms are not fully unveiled, but type-I interferons (TI-IFNs) have been suggested as important mediators of this effect.^9^ Previous research from our group identified a link between the accumulation of TI-IFNs and impairment of T cells to respond to IL-10, a phenomenon associated with the loss of immune tolerance in a model of autoimmunity.^10^

IL-10 is an immunomodulatory molecule that has a non-redundant role in limiting inflammatory responses *in vivo*, particularly in the intestine.^11,12^ This cytokine delivers its functions by acting on a variety of immune cells. It acts on antigen presenting cells decreasing their costimulatory capacity and secretion of inflammatory cytokines,^13,14^ but it also signals directly on T cells.^15^ For example, Tregs need IL-10 stimulation to maintain Foxp3 expression and suppressive capacity^16,17^ and this cytokine also inhibits T cells proliferation and cytokine secretion, modulating T_H_1, T_H_2 and T_H_17 responses.^18–20^ The pathological effects observed when IL-10 signaling in T cells is compromised, underscore the importance of maintaining a functional response on these cells to guarantee the regulation of unwanted reactivity.^15,21^

In the field of transplant tolerance, despite its anticipated protective function, the role of IL-10 remains still controversial. While some reports revealed beneficial effects of exogenous IL-10 signaling in enhancing graft survival, others reported neutral or even detrimental effects.^22–24^ Most of these studies, however, lacked mechanistic insight to understand the variety of outcomes observed after the use of this cytokine as a therapeutic.

In this study, we investigated the role of IL-10 signaling directly in T cells in the therapeutic efficacy of the CoB regimen based on donor specific infusion (DSI) and anti-CD154, in a fully mismatched murine skin allograft model.^25^ We discovered that IL-10 signaling in alloreactive T cells is essential to achieve extension of transplant survival: when disrupted, transplanted mice promptly rejected the allograft despite the CoB treatment. This loss of protection was associated with an increased activation of effector CD4^+^ and CD8^+^ T cells. Interestingly, recapitulating allostimulation *in vitro*, we report that CD4^+^ T cells with impaired IL-10 signaling preserved their ability to differentiate into T_H_1 effectors despite blockade of CD40/CD40L interactions. Moreover, despite integrity of TGF-β signaling, Treg induction and expansion, classically ascribed to the anti-CD154 treatment, was also impaired. Overall, our results show that IL-10 signaling in T cells is essential for CoB regimens and suggest that integrity and optimal engagement of this pathway are necessary for maximization of their therapeutic effect.

## 2. Material and Methods

### 2.1. Mice

BALB/c (H2-K^d^) and C57BL/6 (B6, H2-K^b^) were obtained from the National Cancer Institute (Charles River, Frederick, MD) and IL-10-KO (in B6 background, H2-K^b^) from the Jackson Laboratory. CD4-IL-10R-DN-Foxp3^RFP^-IL10^GFP^ (IL10R-DN) mice were kindly provided by Drs. S. Huber and R.A. Flavell.^15,26^ Animals were housed in specific pathogen-free conditions and the experiments were conducted in accordance with National Institutes of Health guide for use and care of laboratory animals, and under a protocol approved by the JHU Animal Care and Use Committee.

### 2.2. Reagents

Complete medium (CM) consisted of RPMI-1640 (Gibco, Invitrogen, Carlsbad, CA) supplemented as previously described.^10^ Anti-CD154 (clone MR-1) and anti-IL-10R (clone 1B1.3A) monoclonal antibodies were purchased from BioXCell.

### 2.3. Generation of murine bone marrow chimeras

8-week-old B6 mice received two intraperitoneal (IP) doses of Busulfan (MilliporeSigma, 25 mg/Kg) in two consecutive days, followed by intravenous injection, one day after the second dose of busulfan, of 10^7^ million of bone marrow (BM) cells from either B6 or IL10R-DN mice. BM reconstitution in recipient mice was determined via flow cytometry at several time points after injection, with maximum reconstitution achieved 60 days post-procedure **(Supplementary Figure S1).** At that time, BM chimeras were used as recipients in skin transplantation experiments.

### 2.4. T cell and T-depleted splenocytes isolation

For T cell isolation, spleen and lymph nodes were harvested, and total T cells were enriched via magnetic-bead negative selection as previously described.^10^ Briefly, following red blood cell lysis, single cell suspensions were incubated with antibodies anti-CD11b (M1/70), anti-B220 RA3-6B2), anti-Gr-1 (RB6-85C), anti-CD16/32 (2.462), anti-Ter119 (TER-119) and anti-I-A/E (M5/114.15.2 [also anti-CD8 (53-6.7) for CD4 T cell purification, and anti-CD25 (PC61) plus anti-CD44 (IM7) for naïve T cell isolation] (all from BD Biosciences) followed by incubation with anti-rat IgG conjugated magnetic-beads (Thermo Fisher) at 1:1 (cell:bead) ratio. The resulting T cell pool was >90% pure. For T cell-depleted splenocytes, splenic single cell suspensions were incubated with anti-CD3 (17A2), and negative selection performed as described above.

### 2.5. *In vitro* Mix Leucocyte Reaction (MLR) and allogenic Treg differentiation assays

Freshly isolated naïve T cells or naïve CD4^+^ T cells from either B6 or IL-10DN, were stained with cell trace violet (CTV - Thermo Fisher) following manufacturer’s instructions and cocultured for 5 days in U-bottom 96/well plates with BALB/c T-depleted splenocytes at a 1:1 ratio. Cultures were tested with the indicated concentrations of anti-CD154 (MR-1) or TGF-β+IL-2 (both from Peprotech). In all cases, viability, proliferation, cytokine secretion, or Treg percentage at the end of the 5 day-culture were determined by flow cytometry.

### 2.6. Flow cytometry staining

Single cell suspensions from spleen and lymph nodes or recovered from *in vitro* cultures, were either used directly for flow cytometry or stimulated *ex-vivo* for 4h with PMA/ionomycin and cell transport inhibitor (Thermo Fisher) for assessment of intracellular cytokines production. In all studies, cells were first stained with a fixable life/dead probe (Thermo Fisher) to exclude the non-viable population. Thereafter, surface staining was performed with anti-CD3 (17A2), CD4 (RM4-5), CD8 (53-6.7), CD25 (PC61) and CD44 (IM7) (all from Biolegend). When intracellular staining was included, cells were then fixed with the Foxp3 fixation and permeabilization buffer (Thermo Fisher) and subsequently stained with antibodies anti-Foxp3 (FJK-16s, Thermo Fisher), IFN-γ (XMG1.2, BD Biosciences), TNF-α (MP6-XT22, Biolegend), and IL-17A (eBio17B7, Thermo Fisher). Samples were acquired on an LSR-II flow cytometer (BD Biosciences), and results analyzed using FlowJo X version analysis software (FLOWJO LLC, Ashland, OR, USA).

### 2.7. Skin transplantation and immune-modulatory treatments

Full-thickness trunk skin from BALB/c mice was transplanted onto the lateral thoracic wall of B6-background recipients and secured with 6.0 Ethilon sutures.^27^ Skin graft survival was monitored daily. Rejection was defined as 80% necrosis of the skin graft. Where indicated, mice received a regimen consisting of an intravenous (IV) administration of 10^7^ T-depleted splenocytes as Donor Specific Infusion (DSI) on the day of transplantation (post-operative day 0 – POD 0), and 500 μg anti-CD154 (BioXcell), on POD 0 (IV), POD 7 and 14 (Intraperitoneal - IP). Transient pharmacological blockade of IL-10 signaling was achieved by administration of an anti-IL10R antibody (BioXcell), 1mg/week distributed in 6 doses, from POD 0-14.

### 2.8. Western Blot

Naïve and activated CD4^+^ T cells (via 72h incubation with αCD3/28 beads) were either left untreated or stimulated with IL-10 (40 μg/ml, PeproTech) for 20 min and then lysed with RIPA buffer containing proteases and phosphatases inhibitors (MilliporeSigma). Cell lysates were run in 10–12% acrylamide gels and proteins transferred to a PVDF membrane. Protein expression levels were detected with the following antibodies: Jak-1 (6G4) and phospho-Jak-1 (Tyr1034/1035) (from Cell Signaling Technology), IL-10Rα (A3) and β-actin (I-19) (from Santa Cruz Biotechnology). Protein expression was detected using fluorescent-labeled secondary antibodies (LI-COR). Data was acquired using an Odyssey CLx (LI-COR) imaging system.

### 2.9. Statistical analysis

Differences in transplant survival were assessed using a Mantel-Cox log-rank test. Statistical comparison in percentages of T cell viability, proliferation, activation, Treg induction, and secreted cytokines between cell types were determined using unpaired Student’s *t* test. In all cases, results are expressed as mean ± SEM, and analysis were performed with Graph Prism 7 software package (GraphPad, San Diego, CA). A *p*-value <0.05 was considered statistically significant.

## 3. Results

### 3.1. IL-10 signaling is essential for the control of transplant rejection by DSI+anti-CD154

To determine the role of IL-10 signaling in the extension of transplant survival achieved with CoB regimens, we used a full mismatch BALB/c (H-2^d^) to B6 (H-2^b^) skin transplantation model where DSI infusion on POD0 plus a short course of anti-CD154 (on PODs 0, 7 and 14), induces a significant extension of allograft survival when compared to the untreated transplanted group (MST: 102.5 vs 16 d respectively – **Figure 1**). First, we used IL-10-KO mice as recipients. As indicated in Figure 1, IL-10-KO mice showed an almost complete abrogation of the beneficial effect of DSI+anti-CD154 observed in B6-WT mice (MST: 26 vs 102.5 d). Although this outcome clearly relates IL-10 signaling to survival, it is known that the chronic absence of IL-10 in IL-10-KO mice promotes the spontaneous development of inflammatory disease (e.g. colitis),^11^ suggesting that in this model pre-existing conditions could have contributed to the observed accelerated rejection. To exclude this possibility, we transiently blocked IL-10 signaling by treating B6-WT recipients with a blocking antibody anti-IL10R in the period POD 0-14, along with the CoB regimen. This group also showed a significant shortening in graft protection when compared to mice that received CoB only (MST: 47.5 vs 102.5 days - **Figure 1**). These results confirm that IL-10 signaling has an important role in the promotion of transplant survival by CoB.

**Figure 1.**
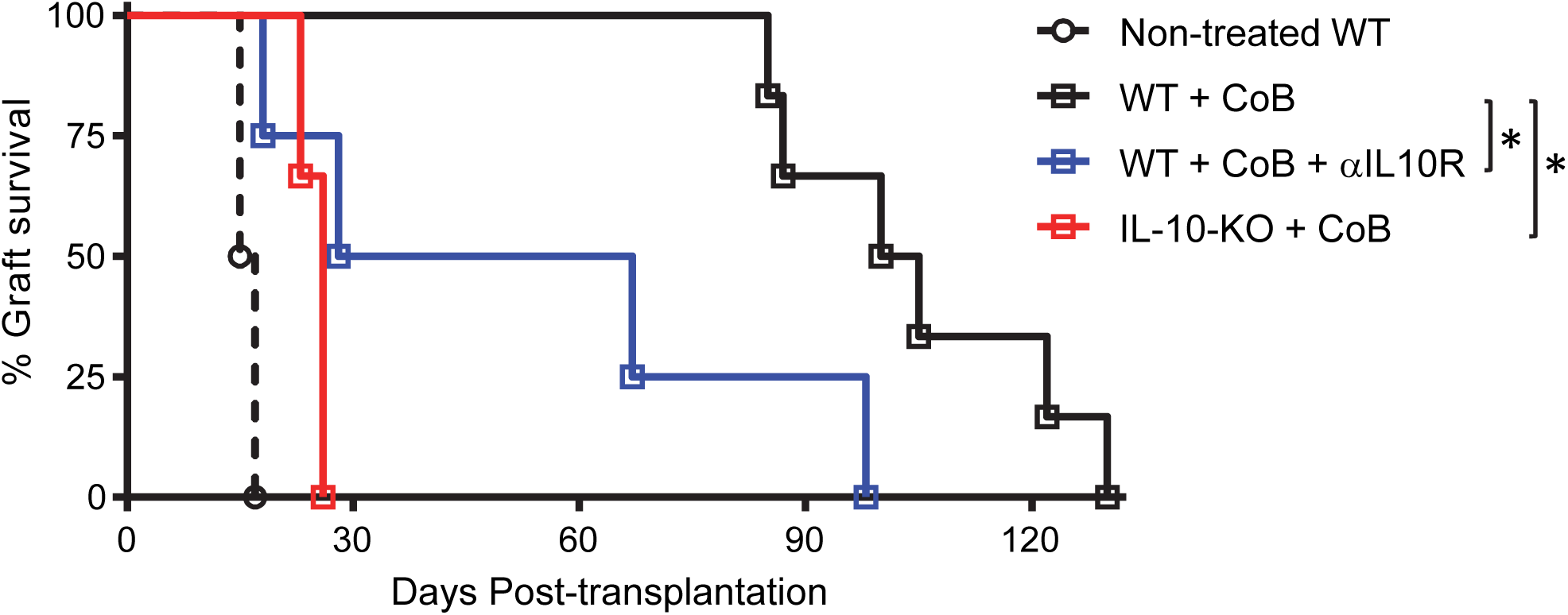
IL-10 signaling is essential for the extension of transplant survival by αCD154+DSI. 8-week-old B6-WT and IL10-KO mice received fully mismatched BALB/c mice skin transplants. Recipients were either left untreated or received a treatment consisting of 10^7^ T-depleted BALB/c splenocytes as Donor Specific Infusion (DSI) intravenously (IV) on POD 0, and 500 μg anti-CD154 (MR-1) on POD 0, 7 and 14 (IV at day 0 and intraperitoneally on day 7 and 14) alone or in combination with 1mg/week of anti-IL10R in 3 doses/week from POD 0-14 (blue line). Graph shows allograft survival curves in B6-WT and IL10-KO treatment groups (n=4-6 animals each). Differences are expressed as *\*p* <0.05, Log-Rank test.

### 3.2. IL-10 signaling in T cells is essential for the efficacy of DSI+anti-CD154

We then focused on understanding the contribution of IL-10 signaling in T cells to graft survival. To this end, we implemented CD4-IL10R-DN mice, where the transgene for a dominant negative version of the IL-10R is controlled by the CD4 promoter, causing impaired IL-10 signaling in all T cells.^15^ First, we created bone marrow (BM) chimeras via reconstitution of B6-WT mice with marrow from either IL-10R-DN mice (IL10R-DN Chimeras) or B6 (WT Chimeras controls). Flow analysis of reconstituted mice indicated that IL10R-DN Chimeras had 30-50% of circulating T cells expressing the transgene (**Supplementary Figure S1**). After receiving skin transplantation and DSI+anti-CD154, the extension of transplant survival in the IL10R-DN-Chimera group was significantly shorter than that of WT-Chimera recipients (MST: 75 vs 142.5 days) (**Figure 2A**). BM-Chimera groups that did not receive CoB treatment rejected the transplants acutely and with overlapping kinetics (**Figure 2A**). We then performed transplants using IL-10R-DN mice as recipients, where all T cells (but only T cells) are impaired in their response to IL-10. In this group, the efficacy of the CoB regimen was significantly reduced relative to the B6-WT group (MST: 29 vs 88 days) (**Figure 2B**). This reduction was comparable to that observed in constitutive IL-10-KO mice (**Figure 1**), suggesting that IL-10 signaling in T cells exerts a dominant role. When comparing the outcome of IL10R-DN-Chimeras and IL-10R-DN recipients (**Figure 2A-B**) a correlation between the percentage of IL-10 non-responsive T cells and CoB efficacy is evident. Similarly to the chimeras experiment, B6 and IL10R-DN transplanted mice that did not received CoB treatment (non-treated), rejected the graft acutely and with similar kinetics (**Figure 2B**).

**Figure 2.**
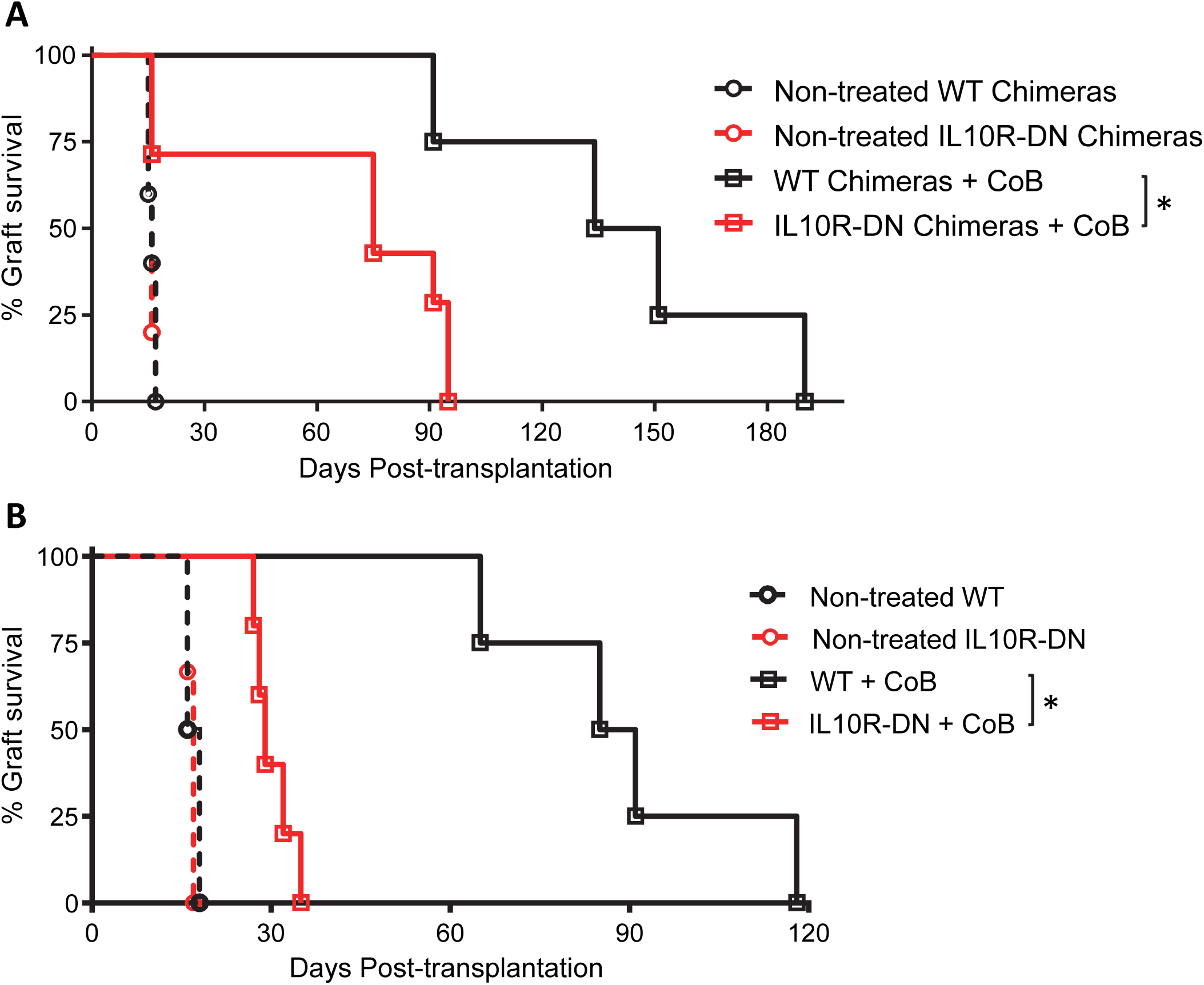
T cells responsiveness to IL-10 is essential for the therapeutic efficacy of αCD154+DSI. **(A)** B6-WT and IL10R-DN-Chimeras (see Material and Methods) or **(B)** WT-B6 and IL10R-DN mice, received fully mismatched BALB/c mice skin transplants. Recipients were either left untreated or received a treatment consisting of 10^7^ T-depleted BALB/c splenocytes (DSI) intravenously (IV) on POD 0, and 500 μg anti-CD154 (MR-1) on POD 0, 7 and 14 (IV at day 0 and intraperitoneally on day 7 and 14). Graph shows allograft survival curves in B6-WT and IL10R-DN-Chimeras groups (n=4-7 animals each) **(A)** and in B6-WT and IL10R-DN recipients (n=4-6 animals each) **(B). (A, B)** Differences are expressed as *\*p* <0.05, Log-Rank test.

### 3.3. CoB treated IL-10R-DN recipient mice present an increased CD4^+^ T cells activation and effector differentiation

Supported by the *in vivo* survival data, we analyzed the phenotype of the T cell compartment associated with the loss of protection in the treated IL10R-DN recipients. We first confirmed that the T cell compartment in unmanipulated mice before transplantation (CD4 and CD8 percentage, T cell effector/memory compartment and Treg percentage) was comparable between IL-10R-DN and their non-transgenic counterparts B6-WT (**Supplementary Figure S2**), as it was also previously reported.^18^ Subsequently, we compared the effector phenotype of T cells in spleen and lymph nodes of non-treated skin recipients at POD 10 after transplantation (**Figure 3**). Both transplanted groups showed a similar increase in CD4^+^ and CD8^+^ effector T cell compartment (CD44^hi^) compared to the non-transplanted controls (**Figure 3B**). The enhanced effector response was associated with an increased secretion of proinflammatory cytokines in both transplanted groups. Percentages of TNF-α and IL-17A secreting CD4^+^ T cells as well as IFN-γ and TNF-α secreting CD8^+^ were higher but comparable in the spleen and lymph nodes of both allograft recipient groups. The proportion of IFN-γ^+^ CD4^+^ T cells was significantly augmented in the spleen of IL10R-DN (**Figure 3C**). Elevated levels of Treg cells were observed in both B6-WT and IL-10R-DN (**Figure 3D**).

**Figure 3.**
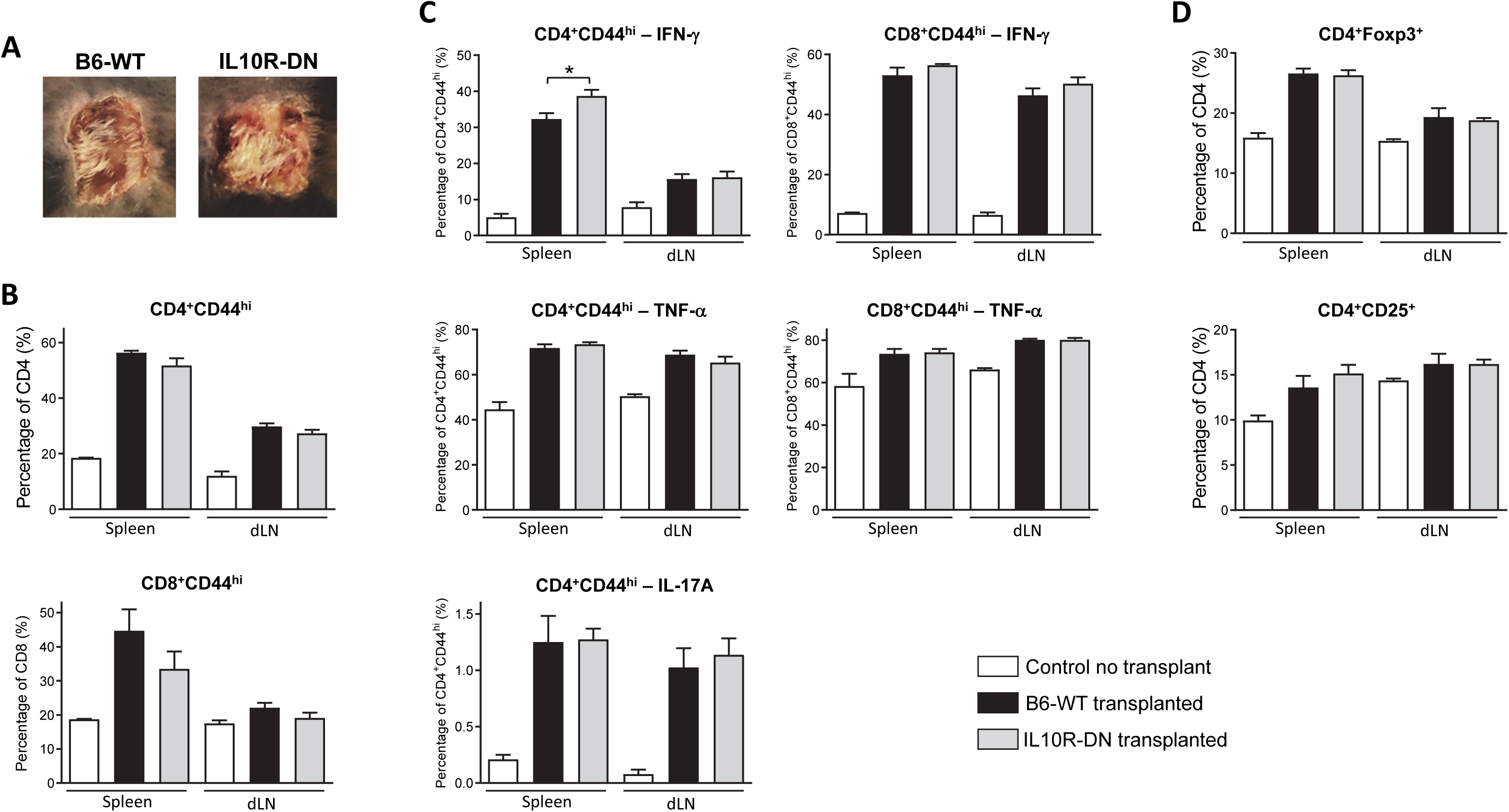
Unmanipulated B6-WT and IL10R-DN recipients have similar T cell alloreactivity. Spleens and draining lymph nodes (axillary ipsilateral to the transplant) were harvested on POD10 from B6-WT and IL10R-DN recipients of BALB/c skin transplants, and the percentages of activated T cells (CD44^hi^), IFN-γ^+^, TNF-α^+^ and IL-17A^+^ CD4/CD8 T cells (post PMA/Ionomycin re-stimulation), as well as Tregs (CD4^+^Foxp3^+^) were characterized by flow cytometry. **(A)** Representative images of the clinical appearance of skin transplants at the moment of sacrifice in each group. **(B-D)** Flow cytometry results showing cumulative results of the percentages of activated (CD44^hi^) CD4/CD8 T cells **(B)**, proinflammatory cytokines (IFN-γ^+^, TNF-α^+^ and IL-17A^+^) producing cells in the CD44^hi^ compartment **(C)**, and Tregs (CD4^+^Foxp3^+^ cells) **(D)**. Data shown in (B-D) are the averages from n=2 independent experiments each generated from 5 individual mice/group and are expressed as percentage ± SEM, *\*p* <0.05 two-tailed unpaired Student’s *t-*test.

We then compared the T cell compartment between CoB-treated B6-WT and IL10R-DN transplanted mice on POD 21 (**Figure 4**). In concordance with the observed extension of transplant survival, there was no significant increase in T cell activation (CD44^hi^ levels in CD4 and CD8 cells) between treated B6-WT recipients and non-transplanted animals (**Figure 4B**). The results revealed, however, a striking expansion of the CD4^+^CD44^hi^ compartment in the IL10R-DN group (**Figure 4B**), despite CoB treatment. The secretion of proinflammatory cytokines followed a similar trend suggesting a lack of control of activation of T cells in the IL10R-DN group. Splenic levels of IFN-γ and IL-17A producing CD4^+^ T cells, as well as of IFN-γ and TNF-α producing CD8^+^ T cells, were significantly higher in the IL10R-DN recipients compared to the B6-WT (**Figure 4C**). Treg cells percentages were augmented in spleen and lymph nodes of both B6-WT and IL10R-DN transplanted mice, but levels reached significance only in B6-WT, when compared to the naive control (**Figure 4D**).

**Figure 4.**
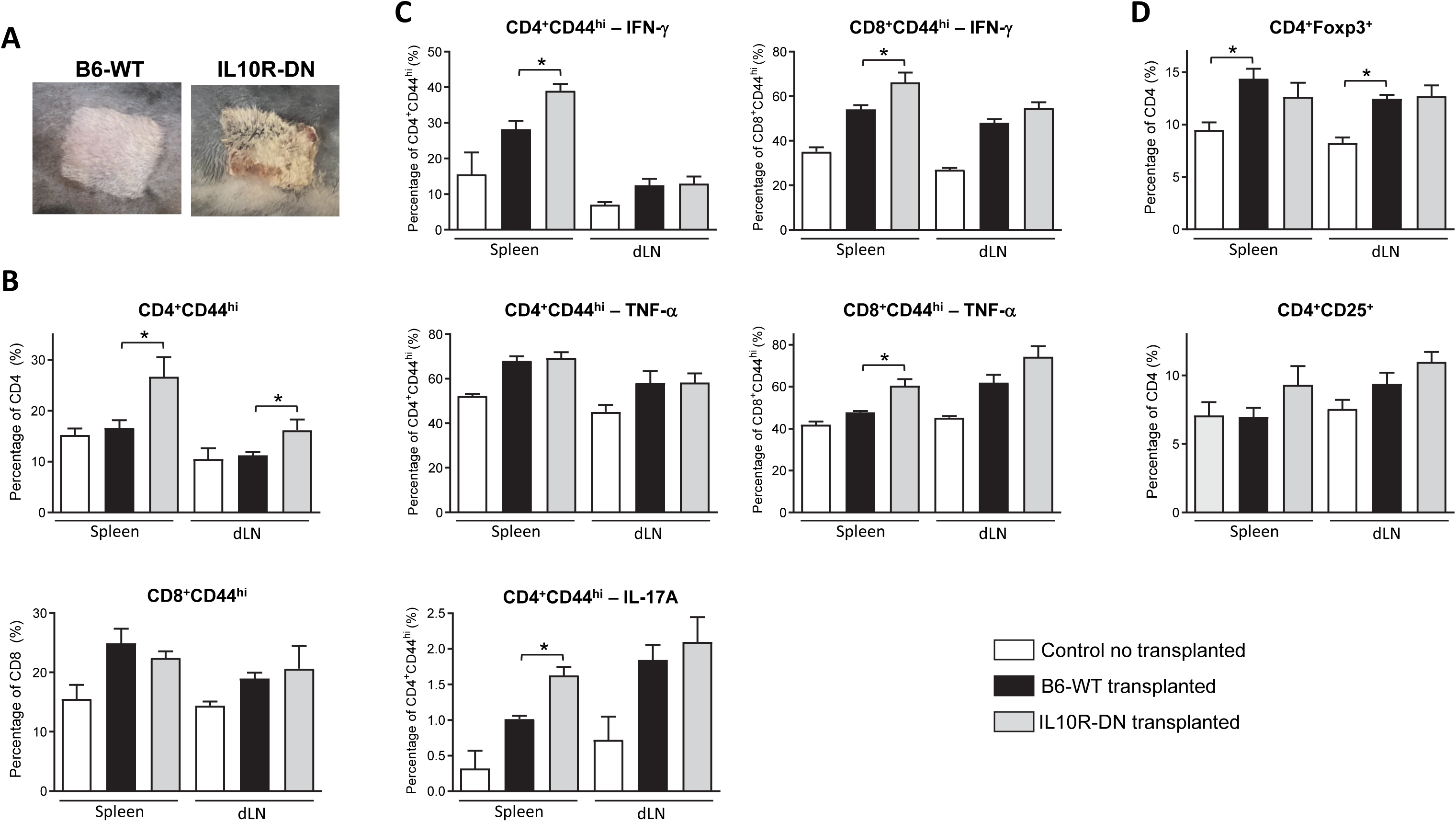
Accelerated rejection in IL10R-DN mice receiving αCD154+DSI is associated with increased T cells activation and differentiation. Spleens and draining lymph nodes (axillary ipsilateral to the transplant) were harvested on POD21 from αCD154+DSI treated B6-WT and IL10R-DN recipients of BALB/c skin transplants, and the percentages of activated T cells (CD44^hi^), IFN-γ^+^, TNF-α^+^ and IL-17A^+^ CD4/CD8 T cells (post PMA/Ionomycin re-stimulation), as well as Tregs (CD4^+^Foxp3^+^) were characterized by flow cytometry. **(A)** Representative images of the clinical appearance of skin transplants at sacrifice in each group. **(B-D)** Flow cytometry results showing cumulative results of the percentage of activated (CD44^hi^) CD4/CD8 T cells **(B)**, proinflammatory cytokines (IFN-γ^+^, TNF-α^+^ and IL-17A^+^) producing cells in the CD44^hi^ compartment **(C)** and Tregs (CD4^+^Foxp3^+^ cells) **(D)**. Data shown in (B-D) are the averages from n=2 independent experiments each generated from 5 individual mice/group and are expressed as percentage ± SEM, *\*p* <0.05 two-tailed unpaired Student’s *t-*test.

### 3.4. CD40-CD154 blockade requires IL-10 signaling in T cells to regulate T_H_1 differentiation and favor Treg induction

The lack of regulation of effector T cell formation in IL-10R-DN recipients suggests that the therapeutic effect of blocking CD40-CD154 interaction depends on direct IL-10 signaling into T cells. To better understand IL-10 function in this context, we set up an MLR assay, using naïve T cells (both CD4 and CD8) from either B6-WT or IL10R-DN mice as responders and allogenic BALB/c T-depleted splenocytes as stimulators. The addition of anti-CD154 mAb did not alter the viability of CD8^+^ T cells at the end of the culture (**Figure 5A**). However, it significantly limited the proliferation and the formation of IFN-γ secreting CD8^+^ T cells (**Figure 5B-C**), as expected. Surprisingly, both B6-WT and IL-10R-DN CD8^+^ T cell responders were similarly affected by CD154 blockade (**Figure 5B-C**). When focusing on the CD4^+^ T cell population, although CD154 blockade reduced proliferation (and not viability) as observed for CD8^+^ T cells (**Figure 6A and B**), the impact on T_H_1 differentiation was different. While IFN-γ secretion was significantly reduced in the B6-WT responders, the proportion of IFN-γ producers remained high in the IL10R-DN group (**Figure 6C**). Analysis via Western Blot of IL-10R levels and IL-10 signaling confirmed that activated CD4^+^ T cells greatly increase their susceptibility and response to this cytokine **(Supplementary Figure S3)**, further confirming the involvement of this signaling pathway in modulating their differentiation in the context of CD154 blockade. In addition to recapitulating T effectors differentiation, our *in vitro* MLR conditions also replicated *in vivo* observations that CD40-CD154 blockade during allostimulation promotes differentiation and expansion of Treg cells. Strikingly, Treg induction was almost completely lost in IL10R-DN responders (**Figure 7A**). This lack of Treg induction was not an intrinsic defect of IL-10R-DN T cells. When alloreactive CD4^+^ T cells were activated in presence of TGF-β, WT and IL-10R-DN cells converted into Foxp3+ iTreg to the same degree and as a direct function of the concentration of TGF-β tested (**Figure 7B**). Altogether, these results demonstrate that induction of IL-10 signaling in T cells is a requirement in the multi-faceted process of immunomodulation exerted by CD154 blockade during activation.

**Figure 5.**
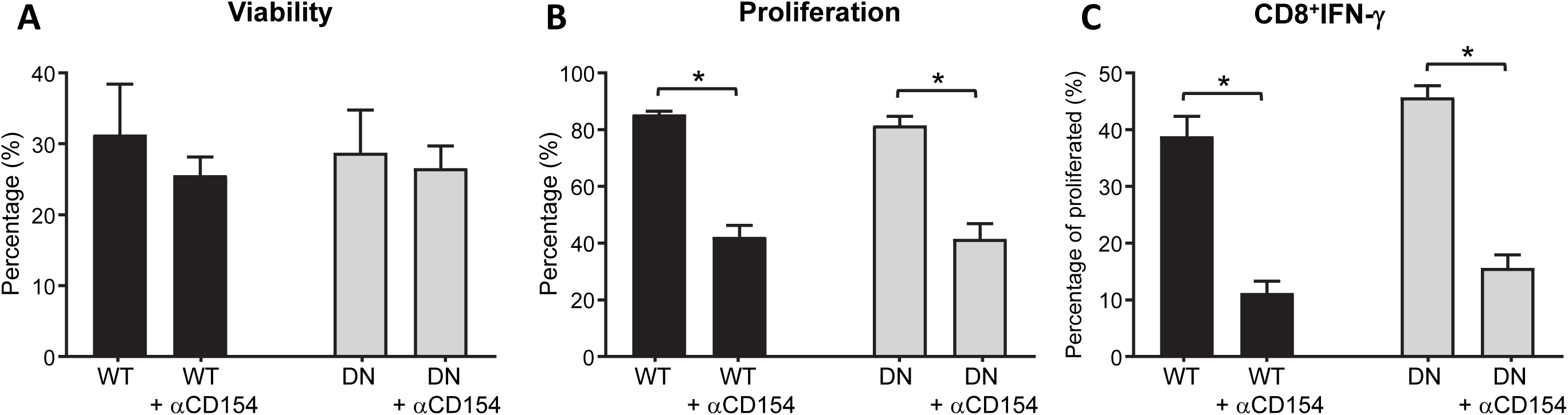
*In vitro* CD154 blockade equally affects proliferation and differentiation of CD8^+^ T cells from B6-WT and IL10R-DN mice. **(A-C)**. CTV-labeled B6-WT or IL10R-DN naïve T cells were cultured for 5 days with allogeneic BALB/c-T depleted splenocytes (1:1, Stimulator to Responder ratio) in presence of anti-CD154 (MR-1, 100 μg/ml). CD8^+^ T cells viability **(A)**, induced proliferation **(B),** and intracellular IFN-γ^+^ levels (post PMA/Ionomycin re-stimulation) in the proliferated population **(C)** were measured by flow cytometry. Data shown is an average of n=3 independent experiments each run in triplicates and expressed as the percentage of viable, proliferated (CTV diluted from the parental peak) or the percentage of IFN-γ secreting within the proliferating CD8^+^ T cells ± SEM with *\*p* <0.05, two-tailed unpaired Student’s *t-*test.

**Figure 6.**
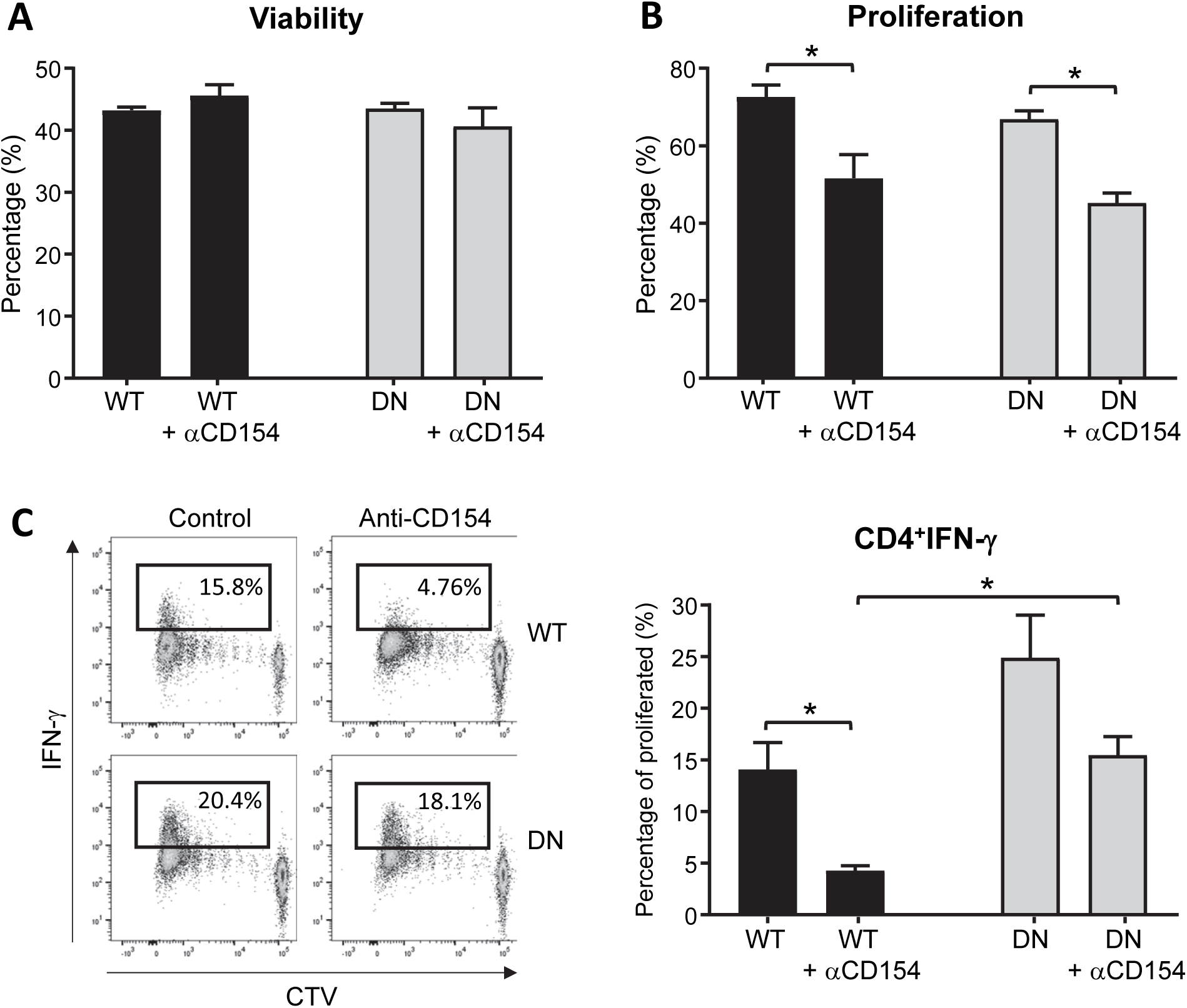
The impact of CD154 blockade on CD4^+^ T cell *in vitro* differentiation is dependent on IL-10 signaling directly in T cells. **(A-C).** CTV-labeled B6-WT or IL10R-DN naïve CD4^+^ T cells were cultured for 5 days with allogeneic BALB/c-T depleted splenocytes (1:1, Stimulator to Responder ratio) in presence of anti-CD154 (MR-1, 100 μg/ml). Viability **(A)**, induced proliferation **(B),** and intracellular IFN-γ^+^ cytokine levels (post PMA/Ionomycin re-stimulation) in the proliferated population **(C)** were measured by flow cytometry. Data shown is a representative (C-left) and average (A-B and C-right) of n=3 independent experiments each run in triplicates and expressed as the percentage of viable, proliferated (CTV diluted from the parental peak) or the percentage of IFN-γ secreting within the proliferating CD4^+^ T cells ± SEM with *\*p* <0.05, two-tailed unpaired Student’s *t-*test.

**Figure 7.**
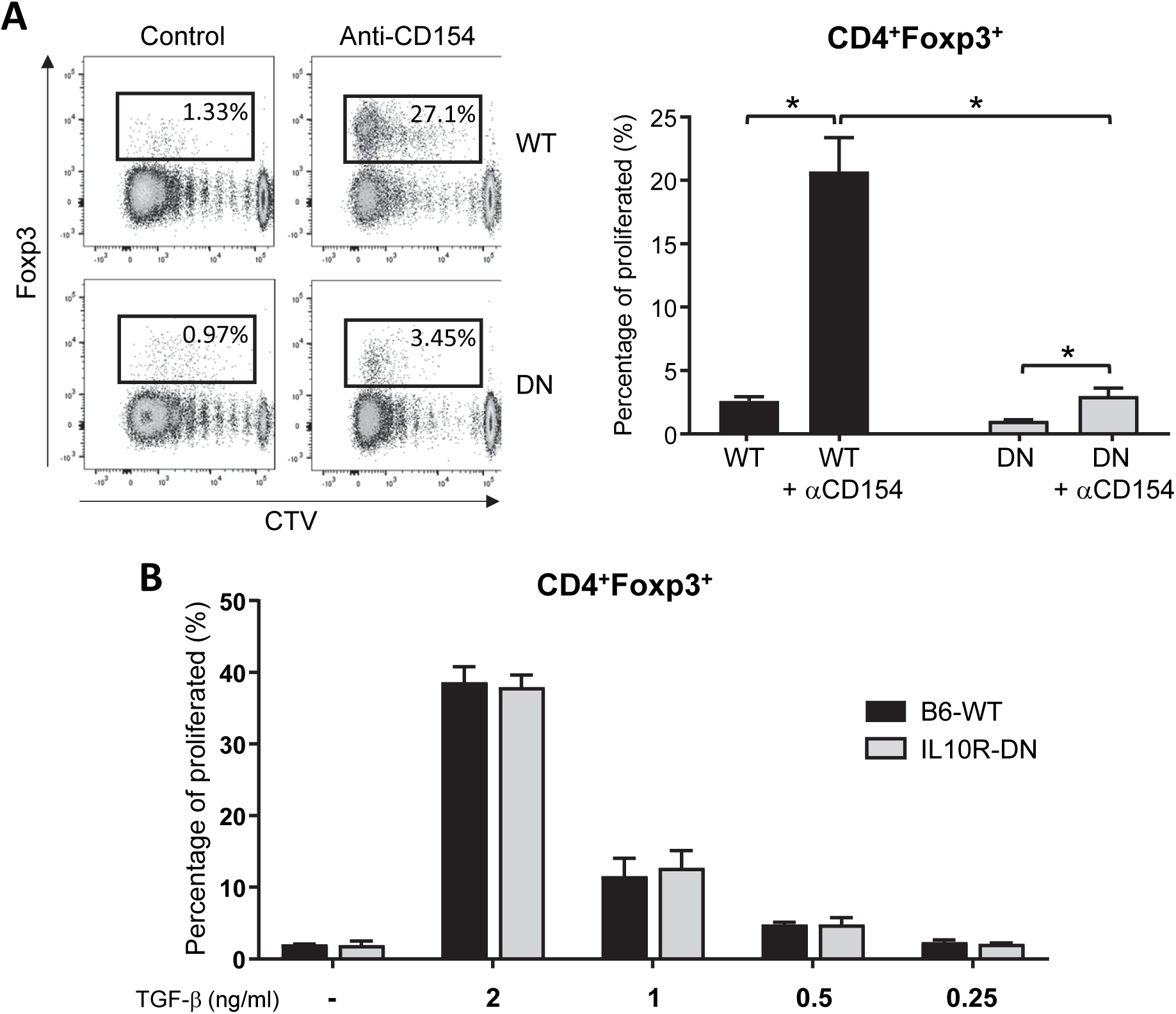
*In vitro* Treg cells induction by CD154 blockade is dependent on IL-10 signaling in T cells. CTV-labeled B6-WT or IL10R-DN naïve CD4^+^ T cells were cultured for 5 days with allogeneic BALB/c-T depleted splenocytes (1:1, Stimulator to Responder ratio) in presence of anti-CD154 (MR-1, 100 μg/ml) alone **(A)** or under Treg inducing conditions (50 U/ml hIL-2 and 0.25 to 2 ng/ml hTGF-β1 titration) **(B)**. The percentage of induced Treg cells (CD4^+^Foxp3^+^) was measured by flow cytometry. Data shown is a representative (A-left) and average (A-right and B) of n=3 independent experiments run in triplicate and expressed as the percentage of Treg cells within the proliferated population (CTV-diluted) comparing B6-WT and IL10R-DN cells at each specific culture condition ± SEM with *\*p* <0.05, two-tailed unpaired Student’s *t-*test.

## 4. Discussion

IL-10 is one of the most important immunomodulatory/anti-inflammatory cytokines, and plays an essential role in the maintenance of immune homeostasis in healthy individuals.^11,12^ However, multiple reports have emphasized that development of therapeutic strategies to exploit the IL-10 response suffers from a still limited understanding of IL-10 biology.^28–32^ Our results provide novel and important insight, as they indicate that IL-10 signaling in T cells is essential to promote the regulation of T_H_1 responses and the induction of Treg cells – hallmarks of blocking the CD40-CD154 pathway for the induction of transplant tolerance.^3,4^ Both our *in vivo* and *in vitro* data show that lack of IL-10 signaling does not intrinsically affect T cells activation (except for an increase in IFN-γ secretion in the CD4 compartment). However, in tolerogenic conditions the outcome of this activation becomes very different when T cells cannot respond to this cytokine.

The capacity of IL-10 to modulate T cell responses is commonly thought to be mainly indirect, through its ability to limit maturation of antigen presenting cells.^13,14^ Our results align with, and add to, recent research indicating that direct signaling of IL-10 on T cells is essential for shaping their activation and differentiation in the context of autoimmunity, cancer, and transplantation.^33^ We observed that in the absence of IL-10 signaling in CD4^+^ T cells, anti-CD154 fails to fully limit the differentiation of IFN-γ secreting cells. IL-10 has been described to limit T_H_1 cells proliferation (and, consequently, IFN-γ production) and IL-2 secretion.^20,34,35^ Recently, the Ziegler’s group suggested a role for this cytokine in countering proliferation and activation of stimulated CD4^+^ T cells by limiting protein synthesis, via inhibition of both mRNA translation and mTORC1 complex activation.^36^ Considering that mTROC1 signaling is a major contributor to the generation of T_H_1 responses,^37^ our results suggest that CD154 blockade could act through modulation of this pathway via IL-10 signaling. Moreover, several reports have highlighted that the transplant survival extension induced by CoB regimens based on anti-CD154 is also due to the induction and expansion of allo-specific Tregs.^4,38,39^ Our findings indicate this effect is completely dependent on the direct signaling of IL-10 in T cells, although the cells still preserve their ability to respond to TGF-β. Important future investigations will need to clarify the molecular mechanisms underlying this prerequisite.

Our in vitro results help us delineate the mechanisms responsible for the limited skin allograft protection induced by CoB in IL10R-DN recipient mice. In addition to secreting this molecule, Tregs need IL-10 stimulation to maintain Foxp3 expression and suppressive capacity,^16,17^ mainly to suppress T_H_17 cells,^17,40,41^ and they are also important regulators of T_H_1 responses.^42^ The increased levels of T_H_1 and T_H_17 cells we observed *in vivo* in CoB treated IL10R-DN recipients could then be promoted by a combination of three factors: i) the lack of the direct effect of IL-10 in modulating CD4^+^ T cells effector differentiation, ii) a compromised Treg induction and expansion, and iii) a lower ability of Tregs to control effector responses. Further studies are needed to confirm whether after CD40/CD154 blockade in allogenic conditions, the absence of IL-10 signaling on T cells also limits the development of other regulatory populations like Tr1 cells,^43,44^ whose induction is IL-10-dependent and can contribute to the induction of allograft tolerance in other settings.^45,46^

The role of IL-10 signaling in CD8^+^ T cells is more controversial, with reports of both stimulatory and inhibitory effects that are probably dependent on the activation context.^23,33^ In our *in vitro* settings, IL-10 signaling in CD8^+^ T cells is dispensable for the inhibitory effect of anti-CD154. However, anti-CD154-treated IL10R-DN transplant recipients show increased accumulation of CD8^+^ secreting proinflammatory cytokines. We hypothesize that the *in vivo* modulation of effector CD8^+^ T cells expansion and function is indirect, through the effect of IL-10 on CD4^+^ T cells (or lack thereof). The discussed lower expansion of Tregs, negative regulators of CD8^+^ T cells activation,^47^ and a sustained licensing of naïve CD8^+^ T cells by activated CD4^+^ T_H_1 cells^48^ could altogether counter the direct effect of anti-CD154 on the cytotoxic T cell compartment.

The blockade of CD40/CD154 interactions, in addition to preventing a full T cell activation and inducing T cell apoptosis or anergy, also favors a tolerogenic DC phenotype, as CD40 engagement in DCs is a mediator of their maturation.^49^ Multiple reports demonstrated that lack of CD40 signaling in DC is associated with inhibition of pro-inflammatory factors secretion and production of IL-10.^50^ Increased levels of IL-10 have been found in the allograft of recipients treated with anti-CD40, and this production has been associated with the expansion of regulatory/tolerogenic myeloid populations.^51^ Interestingly, marginal zone precursor B cells have been suggested to be the main source of IL-10 upon CD154 blockade and the primary driver of allograft tolerance via control of CD4^+^ effector differentiation and induction of regulatory populations.^52,53^ Depletion of marginal zone B cells or their inability to produce IL-10 led to loss of treatment protection and allograft rejection. All these observations suggest that a requirement for the therapeutic activity of CD40-CD154 blockade is the production of IL-10 by antigen presenting cells.

The results presented here showcase the novel concept that the therapeutic efficacy of anti-CD154 is not just a passive effect resulting from the blockade of antigen presenting cells maturation. Instead, it is an active process that requires production and signaling of IL-10 in CD4^+^ T cells (**Figure 8**). It hence highlights the idea that promotion of the effects of IL-10 signaling directly on T cells could be an important point of intervention to limit the negative impact of factors triggering allograft rejection. Considering that inflammatory mediators counter the effect of CoB regimens in transplantation,^6–8^ and our recent observation that TI-IFNs (elevated following inflammatory/infection triggers)^9^ can limit IL-10 signaling in T cells,^10^ our results suggest that identification of a strategy to sustain the effects of IL-10 signaling on T cells would be a valuable approach to enhance the therapeutic efficacy of CoB.^54^ Our results are particularly relevant with the ongoing clinical trials of improved biologics to block the CD40/CD154 axis that have solved the initial thromboembolic complications.^55,56^ For example, we can envision that individual polymorphisms of the IL10/IL10R axis^57^ will probably have a profound influence on the therapeutic outcome of these biologics. Investigations are needed to better delineate the molecular underpinning of the reliance of blocking CD154 on IL-10 signaling. Such understanding will then guide the design of regimens to preserve and amplify IL-10 signaling in T cells (e.g by building upon gene therapy/editing to correct or modify IL-10R expression)^33^ and maximize CoB protocols efficiency toward transplant tolerance.

**Figure 8.**
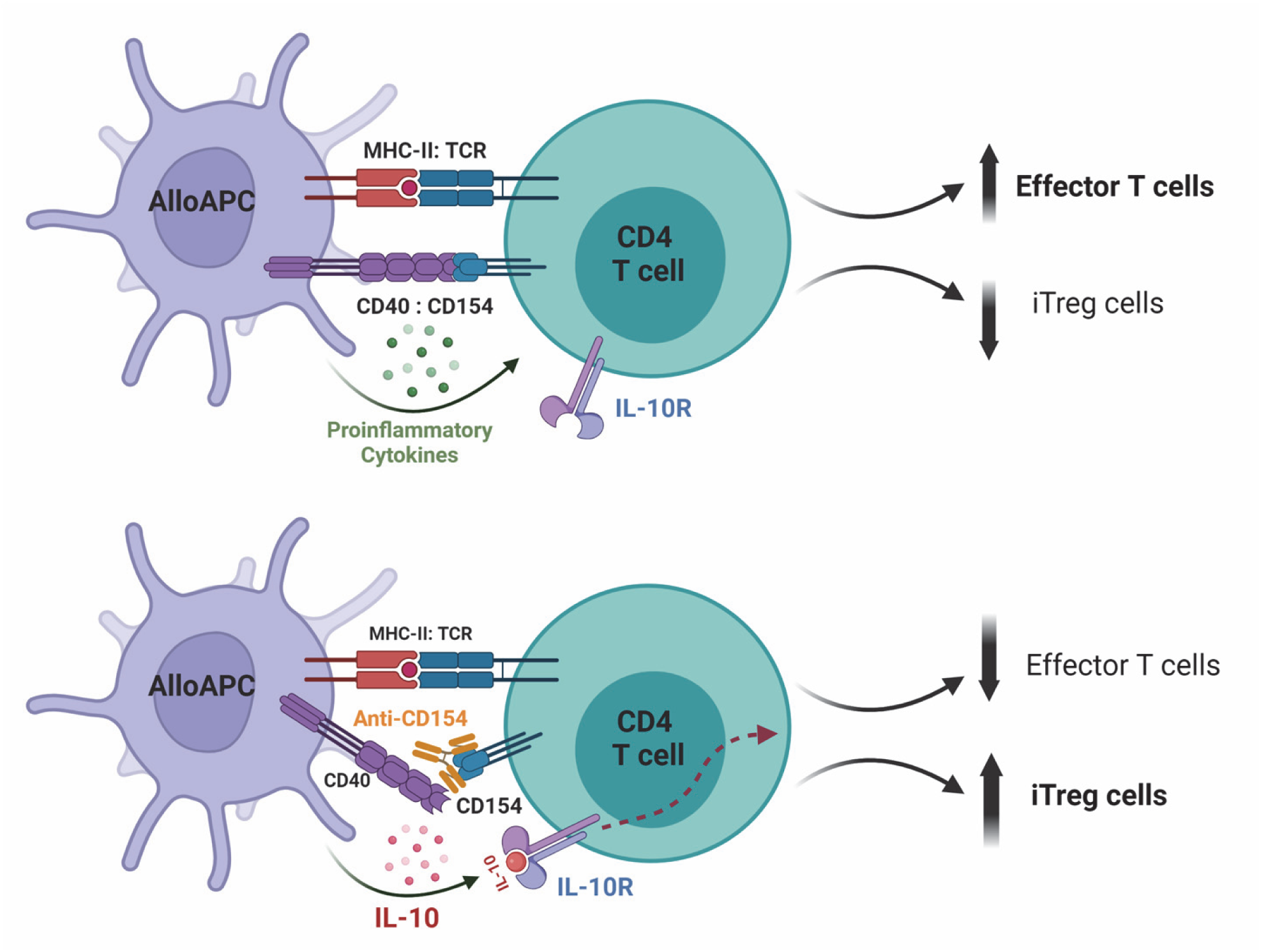
Proposed model of the dependency of CD154 blockade therapeutic efficacy on IL-10 signaling in T cells. <u>Upper Panel</u>, APC engagement of alloreactive CD4^+^ T cells via allogenic MHC-II+peptides presentation and CD40-CD154 interaction promotes CD4^+^ T cell activation, favoring their differentiation into effector populations. <u>Bottom Panel</u>, when presentation occurs in presence of CD154 blockade, IL-10 produced in these conditions acts directly on T cells limiting the differrntation of CD4^+^ T_H_1 effector cells and favoring Treg induction and expansion. Made with BioRender.

B6: C57BL/6 mouse strain
BM: Bone marrow
CM: Complete media
CoB: Costimulation Blockade
DSI: Donor Specific Infusion
IL10R-DN: B6-CD4-IL-10R-DN-Foxp3^RFP^-IL10^GFP^ mouse strain
IP: Intraperitoneal
IV: Intravenous
MLR: Mixed leucocyte reaction
POD: Post-operative day
TI-IFNs: Type-I interferons

## Acknowledgements

This work was supported by NIH grant 1R21HL127355, United States Army Medical Research Acquisition Activity (USMRAA) grant W81XWH-18-1-0789, and internal support from the Dep. of Plastic & Reconstructive Surgery (all to G.R.). The authors thank Samiya Soto and Angela Estevez (laboratory managers), Xiaoling Zhang and Dixie Hoyle (JHU Ross flow cytometry core) for excellent technical assistance and the Division of Research Animal Resources (RAR) at Johns Hopkins University for animal husbandry and care.

## Disclosure

The authors of this manuscript have no conflicts of interest to disclose as described by the *American Journal of Transplantation*.

## Data availability statement

The data that support the findings of this study are available from the corresponding author upon reasonable request.

## Supporting information statement

Additional supporting information may be found online in the Supporting Information section.

## Notes

### Competing Interest Statement

The authors have declared no competing interest.

## References

1. Ford ML, Adams AB, Pearson TC. Targeting co-stimulatory pathways: transplantation and autoimmunity. Nature reviews Nephrology. 2014;10(1):14–24.

2. Yeung MY, Grimmig T, Sayegh MH. Costimulation Blockade in Transplantation. Adv Exp Med Biol. 2019;1189:267–312.

3. Sayegh MH, Turka LA. The role of T-cell costimulatory activation pathways in transplant rejection. N Engl J Med. 1998;338(25):1813–1821.

4. Ferrer IR, Wagener ME, Song M, Kirk AD, Larsen CP, Ford ML. Antigen-specific induced Foxp3+ regulatory T cells are generated following CD40/CD154 blockade. Proc Natl Acad Sci U S A. 2011;108(51):20701–20706.

5. Riella LV, Sayegh MH. T-cell co-stimulatory blockade in transplantation: two steps forward one step back! Expert opinion on biological therapy. 2013;13(11):1557–1568.

6. Thornley TB, Brehm MA, Markees TG, et al. TLR agonists abrogate costimulation blockade-induced prolongation of skin allografts. J Immunol. 2006;176(3):1561–1570.

7. Wang T, Ahmed EB, Chen L, et al. Infection with the intracellular bacterium, Listeria monocytogenes, overrides established tolerance in a mouse cardiac allograft model. Am J Transplant. 2010;10(7):1524–1533.

8. Iglesias M, Khalifian S, Oh BC, et al. A short course of tofacitinib sustains the immunoregulatory effect of CTLA4-Ig in the presence of inflammatory cytokines and promotes long-term survival of murine cardiac allografts. Am J Transplant. 2021;21(8):2675–2687.

9. Thornley TB, Phillips NE, Beaudette-Zlatanova BC, et al. Type 1 IFN mediates cross-talk between innate and adaptive immunity that abrogates transplantation tolerance. J Immunol. 2007;179(10):6620–6629.

10. Iglesias M, Arun A, Chicco M, et al. Type-I Interferons Inhibit Interleukin-10 Signaling and Favor Type 1 Diabetes Development in Nonobese Diabetic Mice. Front Immunol. 2018;9:1565.

11. Kuhn R, Lohler J, Rennick D, Rajewsky K, Muller W. Interleukin-10-deficient mice develop chronic enterocolitis. Cell. 1993;75(2):263–274.

12. Spencer SD, Di Marco F, Hooley J, et al. The orphan receptor CRF2-4 is an essential subunit of the interleukin 10 receptor. J Exp Med. 1998;187(4):571–578.

13. Ding L, Shevach EM. IL-10 inhibits mitogen-induced T cell proliferation by selectively inhibiting macrophage costimulatory function. J Immunol. 1992;148(10):3133–3139.

14. Fiorentino DF, Zlotnik A, Vieira P, et al. IL-10 acts on the antigen-presenting cell to inhibit cytokine production by Th1 cells. J Immunol. 1991;146(10):3444–3451.

15. Kamanaka M, Huber S, Zenewicz LA, et al. Memory/effector (CD45RB(lo)) CD4 T cells are controlled directly by IL-10 and cause IL-22-dependent intestinal pathology. J Exp Med. 2011;208(5):1027–1040.

16. Murai M, Turovskaya O, Kim G, et al. Interleukin 10 acts on regulatory T cells to maintain expression of the transcription factor Foxp3 and suppressive function in mice with colitis. Nat Immunol. 2009;10(11):1178–1184.

17. Chaudhry A, Samstein RM, Treuting P, et al. Interleukin-10 signaling in regulatory T cells is required for suppression of Th17 cell-mediated inflammation. Immunity. 2011;34(4):566–578.

18. Huber S, Gagliani N, Esplugues E, et al. Th17 cells express interleukin-10 receptor and are controlled by Foxp3(-) and Foxp3+ regulatory CD4+ T cells in an interleukin-10-dependent manner. Immunity. 2011;34(4):554–565.

19. Coomes SM, Kannan Y, Pelly VS, et al. CD4(+) Th2 cells are directly regulated by IL-10 during allergic airway inflammation. Mucosal Immunol. 2017;10(1):150–161.

20. Taga K, Tosato G. IL-10 inhibits human T cell proliferation and IL-2 production. J Immunol. 1992;148(4):1143–1148.

21. Iyer SS, Cheng G. Role of interleukin 10 transcriptional regulation in inflammation and autoimmune disease. Critical reviews in immunology. 2012;32(1):23–63.

22. Bromberg JS. IL-10 immunosuppression in transplantation. Curr Opin Immunol. 1995;7(5):639–643.

23. Moore KW, de Waal Malefyt R, Coffman RL, O’Garra A. Interleukin-10 and the interleukin-10 receptor. Annu Rev Immunol. 2001;19:683–765.

24. Kim YH, Lim DG, Wee YM, et al. Viral IL-10 gene transfer prolongs rat islet allograft survival. Cell Transplant. 2008;17(6):609–618.

25. Wang T, Chen L, Ahmed E, et al. Prevention of allograft tolerance by bacterial infection with Listeria monocytogenes. J Immunol. 2008;180(9):5991–5999.

26. Kamanaka M, Kim ST, Wan YY, et al. Expression of interleukin-10 in intestinal lymphocytes detected by an interleukin-10 reporter knockin tiger mouse. Immunity. 2006;25(6):941–952.

27. Cheng CH, Lee CF, Fryer M, et al. Murine Full-thickness Skin Transplantation. J Vis Exp. 2017(119).

28. Couper KN, Blount DG, Riley EM. IL-10: the master regulator of immunity to infection. J Immunol. 2008;180(9):5771–5777.

29. Shinozaki K, Yahata H, Tanji H, Sakaguchi T, Ito H, Dohi K. Allograft transduction of IL-10 prolongs survival following orthotopic liver transplantation. Gene Ther. 1999;6(5):816–822.

30. Xiong J, Wang Y, Zhang Y, et al. Lack of Association between Interleukin-10 Gene Polymorphisms and Graft Rejection Risk in Kidney Transplantation Recipients: A Meta-Analysis. PLoS One. 2015;10(6):e0127540.

31. Blazar BR, Taylor PA, Smith S, Vallera DA. Interleukin-10 administration decreases survival in murine recipients of major histocompatibility complex disparate donor bone marrow grafts. Blood. 1995;85(3):842–851.

32. Bedke T, Muscate F, Soukou S, Gagliani N, Huber S. Title: IL-10-producing T cells and their dual functions. Semin Immunol. 2019;44:101335.

33. Saraiva M, Vieira P, O’Garra A. Biology and therapeutic potential of interleukin-10. J Exp Med. 2020;217(1).

34. Taylor A, Verhagen J, Akkoc T, et al. IL-10 suppresses CD2-mediated T cell activation via SHP-1. Mol Immunol. 2009;46(4):622–629.

35. de Waal Malefyt R, Figdor CG, Huijbens R, et al. Effects of IL-13 on phenotype, cytokine production, and cytotoxic function of human monocytes. Comparison with IL-4 and modulation by IFN-gamma or IL-10. J Immunol. 1993;151(11):6370–6381.

36. So L, Obata-Ninomiya K, Hu A, et al. Regulatory T cells suppress CD4+ effector T cell activation by controlling protein synthesis. J Exp Med. 2023;220(3).

37. Delgoffe GM, Pollizzi KN, Waickman AT, et al. The kinase mTOR regulates the differentiation of helper T cells through the selective activation of signaling by mTORC1 and mTORC2. Nat Immunol. 2011;12(4):295–303.

38. Burrell BE, Bromberg JS. Fates of CD4+ T cells in a tolerant environment depend on timing and place of antigen exposure. Am J Transplant. 2012;12(3):576–589.

39. Pinelli DF, Wagener ME, Liu D, et al. An anti-CD154 domain antibody prolongs graft survival and induces Foxp3(+) iTreg in the absence and presence of CTLA-4 Ig. Am J Transplant. 2013;13(11):3021–3030.

40. Diefenhardt P, Nosko A, Kluger MA, et al. IL-10 Receptor Signaling Empowers Regulatory T Cells to Control Th17 Responses and Protect from GN. J Am Soc Nephrol. 2018;29(7):1825–1837.

41. Hara M, Kingsley CI, Niimi M, et al. IL-10 is required for regulatory T cells to mediate tolerance to alloantigens in vivo. J Immunol. 2001;166(6):3789–3796.

42. Koch MA, Tucker-Heard G, Perdue NR, Killebrew JR, Urdahl KB, Campbell DJ. The transcription factor T-bet controls regulatory T cell homeostasis and function during type 1 inflammation. Nat Immunol. 2009;10(6):595–602.

43. Gregori S, Tomasoni D, Pacciani V, et al. Differentiation of type 1 T regulatory cells (Tr1) by tolerogenic DC-10 requires the IL-10-dependent ILT4/HLA-G pathway. Blood. 2010;116(6):935–944.

44. Brockmann L, Gagliani N, Steglich B, et al. IL-10 Receptor Signaling Is Essential for TR1 Cell Function In Vivo. J Immunol. 2017;198(3):1130–1141.

45. Fousteri G, Jofra T, Di Fonte R, et al. Lack of the protein tyrosine phosphatase PTPN22 strengthens transplant tolerance to pancreatic islets in mice. Diabetologia. 2015;58(6):1319–1328.

46. Gagliani N, Jofra T, Stabilini A, et al. Antigen-specific dependence of Tr1-cell therapy in preclinical models of islet transplant. Diabetes. 2010;59(2):433–439.

47. McNally A, Hill GR, Sparwasser T, Thomas R, Steptoe RJ. CD4+CD25+ regulatory T cells control CD8+ T-cell effector differentiation by modulating IL-2 homeostasis. Proc Natl Acad Sci U S A. 2011;108(18):7529–7534.

48. Tuzlak S, Dejean AS, Iannacone M, et al. Repositioning T(H) cell polarization from single cytokines to complex help. Nat Immunol. 2021;22(10):1210–1217.

49. Quezada SA, Jarvinen LZ, Lind EF, Noelle RJ. CD40/CD154 interactions at the interface of tolerance and immunity. Annu Rev Immunol. 2004;22:307–328.

50. Tang T, Cheng X, Truong B, Sun L, Yang X, Wang H. Molecular basis and therapeutic implications of CD40/CD40L immune checkpoint. Pharmacol Ther. 2021;219:107709.

51. Conde P, Rodriguez M, van der Touw W, et al. DC-SIGN(+) Macrophages Control the Induction of Transplantation Tolerance. Immunity. 2015;42(6):1143–1158.

52. Lal G, Kulkarni N, Nakayama Y, et al. IL-10 from marginal zone precursor B cells controls the differentiation of Th17, Tfh and Tfr cells in transplantation tolerance. Immunology letters. 2016;170:52–63.

53. Lal G, Nakayama Y, Sethi A, et al. Interleukin-10 From Marginal Zone Precursor B-Cell Subset Is Required for Costimulatory Blockade-Induced Transplantation Tolerance. Transplantation. 2015;99(9):1817–1828.

54. Mori DN, Kreisel D, Fullerton JN, Gilroy DW, Goldstein DR. Inflammatory triggers of acute rejection of organ allografts. Immunol Rev. 2014;258(1):132–144.

55. Lassiter G, Otsuka R, Hirose T, et al. TNX-1500, a crystallizable fragment-modified anti-CD154 antibody, prolongs nonhuman primate renal allograft survival. Am J Transplant. 2023;23(8):1171–1181.

56. Miura S, Habibabady ZA, Pollok F, et al. TNX-1500, a crystallizable fragment-modified anti-CD154 antibody, prolongs nonhuman primate cardiac allograft survival. Am J Transplant. 2023;23(8):1182–1193.

57. Turner DM, Williams DM, Sankaran D, Lazarus M, Sinnott PJ, Hutchinson IV. An investigation of polymorphism in the interleukin-10 gene promoter. Eur J Immunogenet. 1997;24(1):1–8.

