## Supplementary Data for "T cell responsiveness to IL-10 defines the immunomodulatory effect of costimulation blockade via anti-CD154 and impacts transplant survival"

### Supporting information

#### Supplementary Figure S1

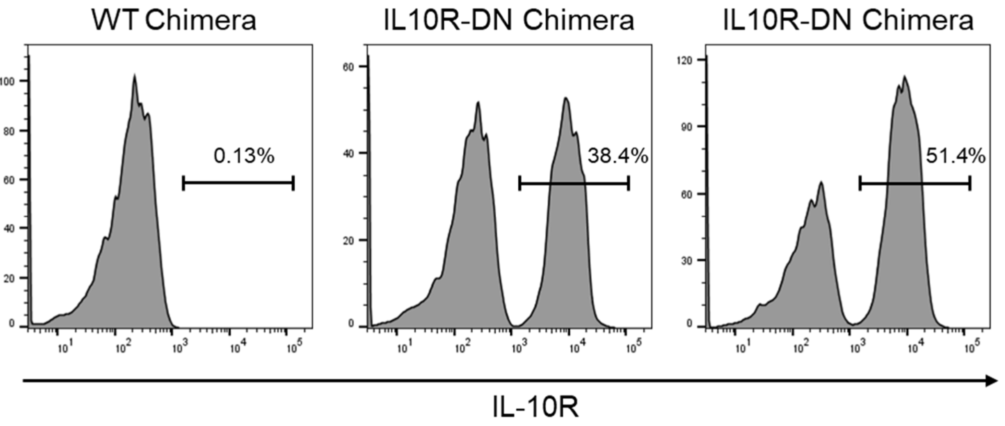


**Supplementary Figure S1. Chimerism levels at the time of skin transplantation in mixed bone marrow chimeras.** The percentage of IL10R-DN expressing T cells in B6 mice infused with bone marrow from either B6-WT or IL10R-DN donors (WT Chimeras and IL10R-DN Chimeras respectively) was measured in blood samples by flow cytometry, 60 days post-bone marrow infusion. Data shown are representative of individual animals and are expressed as the percentage of IL10R$^+$ cells within the CD3 population.

**Supplementary Figure S2**

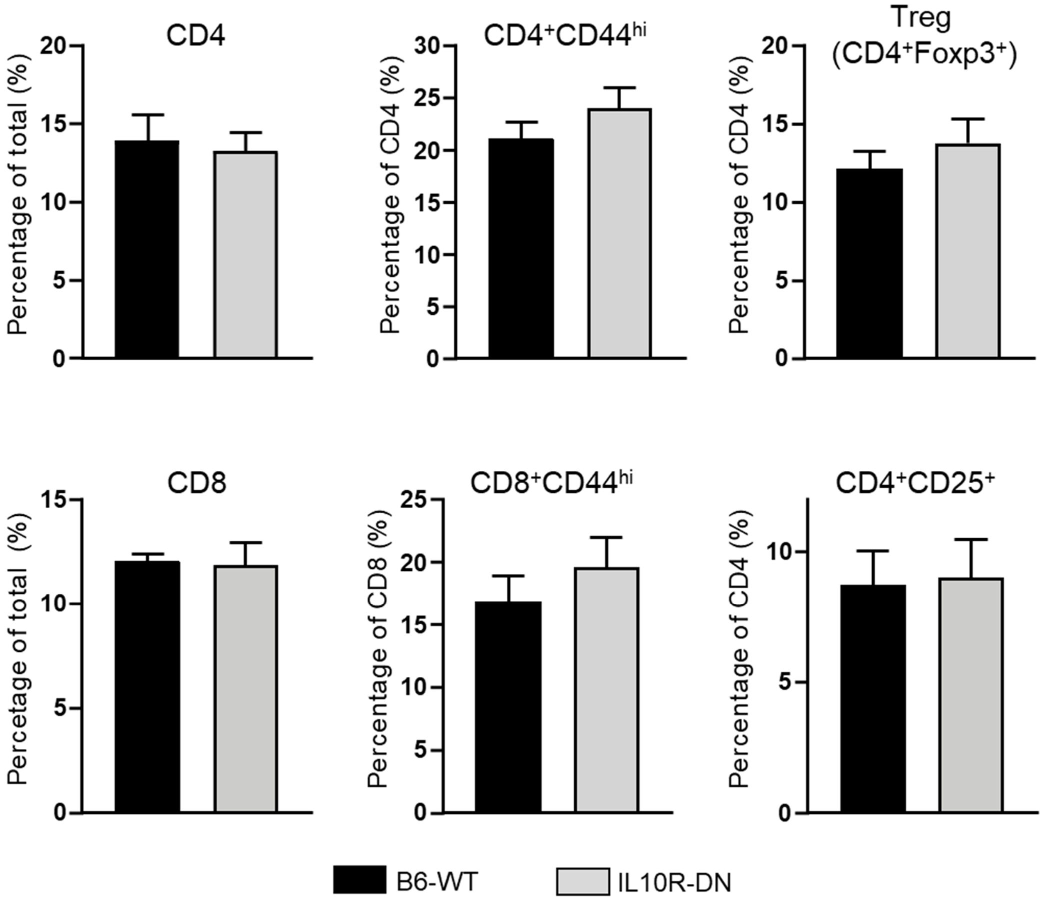


**Supplementary Figure S2. Comparison of T cell compartments between unmanipulated B6-WT and IL10R-DN mice pre-transplantation.** Spleens were harvested from 8-10 weeks old naïve B6 and IL-10R-DN mice and the percentage of CD4, CD8, $CD44^{hi}$, $CD25^{+}$ and $Foxp3^{+}$ cells were measured by flow cytometry. Data shown is an average of n=4 independent mice per group and expressed as the percentage of cells within the total or specific subpopulations ± SEM with *$p$ <0.05, two-tailed unpaired Student's *t*-test.

**Supplementary Figure S3**

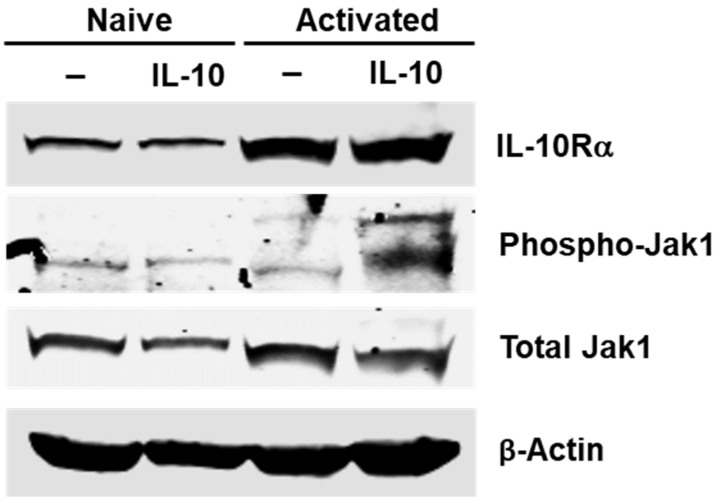


**Supplementary Figure S3. Response to IL-10 in Naïve and Activated CD4+ T cells.** Naïve and activated CD4+ T cells (via 72h stimulation with αCD3/28-beads) were either left untreated or stimulated with IL-10 (40 µg/ml) for 20 min. Protein expression of molecules involved in IL-10 signaling (IL10Rα, Jak-1, phospho-Jak-1) was measured in each condition by western blot. Detection of β-actin was used as a housekeeping loading control.
